# Machine learning predicts system-wide metabolic flux control in cyanobacteria

**DOI:** 10.1101/2023.10.18.562898

**Authors:** Amit Kugler, Karin Stensjö

## Abstract

Metabolic fluxes and their control mechanisms are fundamental in cellular metabolism, offering insights for the study of biological systems and biotechnological applications. However, quantitative and predictive understanding of controlling biochemical reactions in microbial cell factories, especially at the system level, is limited. In this work, we present ARCTICA, a computational framework that integrates constraint-based modelling with machine learning tools to address this challenge. Using the model cyanobacterium *Synechocystis* sp. PCC 6803 as chassis, we demonstrate that ARCTICA effectively simulates global-scale metabolic flux control. Key findings are that (i) the photosynthetic bioproduction is mainly governed by enzymes from the Calvin–Benson–Bassham (CBB) cycle, rather than by those involve in the biosynthesis of the end-product, (ii) the catalytic capacity of the CBB cycle limits the photosynthetic activity and downstream pathways and (iii) ribulose-1,5-bisphosphate carboxylase/oxygenase (RuBisCO) is a major, but not the most, limiting step in the CBB cycle. Predicted metabolic reactions qualitatively align with prior experimental observations, validating our modelling approach. ARCTICA serves as a valuable pipeline for understanding cellular physiology and predicting rate-limiting steps in genome-scale metabolic networks, providing guidance for bioengineering of cyanobacteria.

**Highlights:**

1. A workflow for flux control analysis in *Synechocystis* sp. PCC 6803.
2. Machine learning with features derived from genome-scale metabolic modelling.
3. Identification of potential key reactions for metabolic adaptations and cell bioengineering.

## Introduction

Photoautotrophic microorganisms, such as cyanobacteria, are regarded as promising platforms for the sustainable manufacturing of various chemicals, based on their inherent ability to absorb light and fix CO2, thereby producing organic molecules through a photo-driven process (Kukil & Lindberg, 2022; Liu et al., 2019; Mustila et al., 2021; Rodrigues & Lindberg, 2021). Yet, the light-to-product conversion efficiency has be to be improved in these cell factories in order to fully exploit their potential.

Several efforts to identify limiting steps in cellular metabolism have been recorded. Main approaches being employed consist of ^13^C-based metabolic flux analysis (MFA) (Sauer, 2004) and metabolic control analysis (MCA) using kinetic models (Heinrich & Rapoport, 1973; Kacser & Burns, 1973). Fluxomic studies assisted in the improvement in isobutyraldehyde and 2,3-butanediol production rate in *Synechococcus elongatus* PCC 7942 (Jazmin et al., 2017; Kanno et al., 2017). Kinetic models, though at local scale, have also been used to understand cyanobacterial metabolism (Jablonsky et al., 2013, 2016; Janasch et al., 2018), and to identify overexpression targets for enhanced production of limonene (Wang et al., 2016) and ethanol (Nishiguchi et al., 2019). While MFA offers precise and detailed quantification of the intracellular fluxes in the living cell, the high experimental demand of this technique hinders the wide application for systematic metabolic engineering analyses (Sauer, 2004). The construction of kinetic models is challenging too due the required knowledge about kinetic parameter values (e.g., Michaelis constants), which are scarce (Davidi & Milo, 2017; Islam et al., 2021; Nilsson et al., 2017), especially for cyanobacteria (Chang et al., 2021; Wittig et al., 2018).

In contrast, constraint-based analysis (CBA) uses linear programming, which allows for metabolic modelling at the genome scale, by setting reaction constraints and an objective function (Orth et al., 2010). Thereafter, in silico flux distributions through the biochemical network can be calculated upon optimization of a chosen objective (e.g., maximal growth rate). Since the number of reactions in the metabolic network exceeds the number of metabolites, the system is said to be underdetermined. On the one hand, this could lead to overprediction of metabolic capabilities, for which the solution space can be minimized by integrating quantitative omics data into the reconstructed model (Reed, 2012). On the other hand, this inherent property provides an opportunity to study the robustness of the metabolic reactions within the network, which can be realized through, for example, protein and metabolic engineering (Greenhalgh et al., 2021).

Earlier studies have proposed CBA-based methods for selecting targets for engineering (Choi et al., 2010; Ranganathan et al., 2010; Tsouka et al., 2021). These analyses maximize a singular objective function and are thus biased towards the investigated bioproduct. In reality, however, the cell has multiple objectives to satisfy. Monte-Carlo sampling method, on the contrary, generates an unbiased probability of the steady-state flux distributions, which are solely based on the network topology and the imposed constraints (Schellenberger & Palsson, 2009; Wiback et al., 2004). Such technique was proven useful for revealing transcriptional regulation in key enzymes (Bordel et al., 2010), differentiating between strains producing specific compounds (Scott et al., 2021), as well as for deciphering the effect of environmental conditions on the flux distributions (Herrmann et al., 2019; Pearcy et al., 2022; Wanichthanarak et al., 2020).

In recent years, an integrative approach, combining machine learning (ML) and CBA, has become a key element in addressing biological questions. Results derived from CBA simulations were processed by ML models, which improved the interpretability of the CBA results. Examples include the classification of metabolic fluxes based on drug side effects (Shaked et al., 2016), as well as identification of growth-limiting factors from flux distribution (Vijayakumar et al., 2020). Alternatively, ML algorithms were used to improve the predictive capability of CBA, by integrating enzyme biochemical characteristics (Heckmann et al., 2018, 2020) or other experimental data (Zhang et al., 2020).

Here, we present ARCTICA (mAchine leaRning for Constrained-based meTabolIc Control Analysis) to identify key controlling reactions that influence bioproduction at the genome-scale, based on the changes in internal fluxes. Using the model cyanobacterium *Synechocystis* sp. PCC 6803 (hereafter *Synechocystis*) as chassis, we illustrate the utility of ARCTICA for assessing the metabolism of cyanobacteria producing various food ingredients, as well as discuss its advantages and limitations.

## Methods

### Computational tools

Flux balance analysis (Orth et al., 2010) was employed on COBRApy v0.20.035 using the commercial solver Gurobi v10.0.1 (Gurobi Optimization, Inc., Houston, TX, United States) in Python v3.10.12, using the packages Pandas v1.5.3, NumPy v1.23.5, Matplotlib v3.7.1, Seaborn v0.12.2 and Pickle v4.0. Machine learning algorithm was applied using scikit-learn v1.2.2 (Pedregosa et al., 2011). Visualization of the biological pathways was done using the Escher web-tool (King et al., 2015).

### Metabolic network reconstruction

The iJN678_AK genome-scale metabolic model of *Synechocystis* was used for the computational analysis (Kugler & Stensjö, 2023). For the simulation of recombinant *Synechocystis* producing metabolites of interest, the metabolic reconstruction was expanded with unidirectional pseudo-reactions allowing product export of the respective compounds (Supplementary Tables 1-3). For lauric acid production, a heterologous thioesterase reaction was added, allowing the conversion of lauryl-ACP to lauric acid (Supplementary Table 1).

### Constraint-based modelling

To evaluate the maximum production rate of end-products, flux balance analysis was employed, using COBRApy (Ebrahim et al., 2013). To estimate the flux solution space, Monte-Carlo flux sampling (Schellenberger & Palsson, 2009; Wiback et al., 2004) was carried out with the optGpSampler method (Megchelenbrink et al., 2014) implemented in COBRApy, and using 100,000 sample points of randomly distributed fluxes, and a thinning factor of 100. Prior to sampling, the exchange reaction rate for metabolite excretion was set to 10% and 90% of the predicted maximum production rate at each run. Then, to capture the metabolic mechanisms underlying responses to perturbations, the obtained flux values were normalized to the export rate of the end-product, providing percentage of flux values in the metabolic pathways.

### Machine learning and feature selection

The multiple datasets generated by flux sampling were concatenated into a single comprehensive data matrix (200,000 × 885), to which a machine learning algorithm was applied. To ensure model generalization, we randomly split the samples dataset into train and test subsets, comprising 67% and 33% of the main dataset, respectively, with ten-fold cross-validation. To identify top-contributing features (fluxomic predictors), we used logistic regression model with an L1 penalty (Least Absolute Shrinkage and Selection Operator; LASSO) (Tibshirani, 1996). The predictive accuracy of the ML model was evaluated by assessing the accuracy scores and the classification report. For clarity, exchange and transport reactions (either naturally occurring or necessary for the modelling) were omitted from the analysis.

## Results

### Construction of ARCTICA for predicting important metabolic reactions

In order to extract and rank rate-controlling steps, that upon fine-tuning would potentially enhance the production rate of target molecules, we developed a computational framework (ARCTICA), integrating constraint-based metabolic modelling with machine learning tools (Figure 1). We leverage the undetermined system of flux balance analysis, resulting in alternative internal flux distributions, and performed MCA at the genome scale, based on reaction rates. The rationale behind this is that metabolic fluxes are the ultimate output of enzyme catalysis, forming the basis for cellular operation (Nielsen, 2003; Sauer, 2006). Thence, understanding how an alteration in the reaction rates themselves would affect the flux of the other reactions in the network would be the ideal analysis of metabolic control.

**Figure 1.**
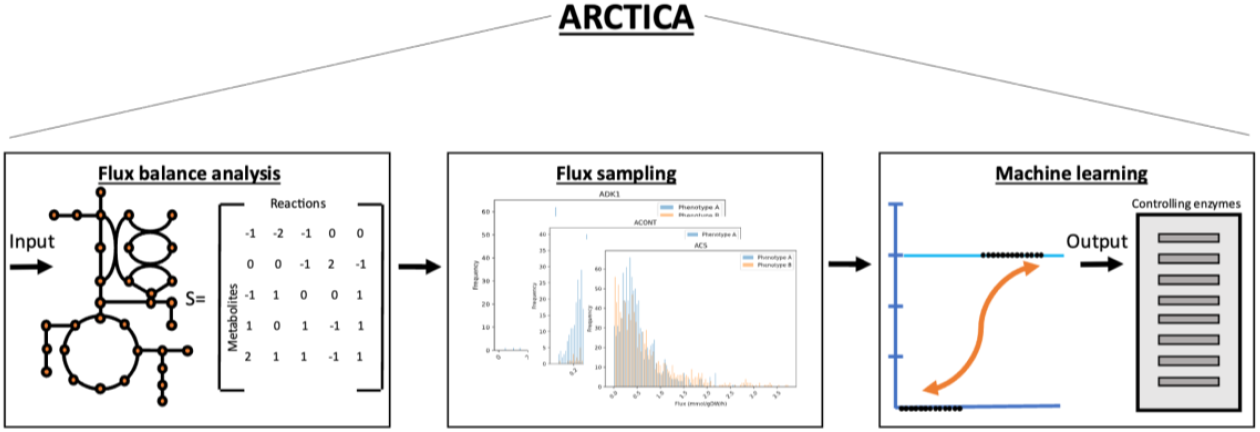
Flowchart of the computational framework (ARCTICA) proposed in this study for predicting metabolic flux control. The input for ARCTICA is defined by setting the light and carbon uptake rates.

ARCTICA consists of four steps: (i) Implementation of flux balance analysis to compute the maximal production of the metabolite of interest. (ii) Constraining the excretion rate of the selected metabolite to 10% and 90% of maximal production rate to yield two in silico phenotypes (high-producing and low-producing strains). (iii) Application of Monte Carlo random flux sampling to compute the probability for feasible fluxes that satisfy the network constraints. (iv) Generation of a comprehensive data matrix by concatenating the obtained datasets to be employed as features (metabolic fluxes) for training the machine learning model. Thereafter, using logistic regression and feature selection, metabolic reactions that statistically influence and distinguish the two phenotypes can be mined.

For evaluating the applicability of ARCTICA, we examined the network reconstruction of the model cyanobacterium *Synechocystis*, and chose three metabolites which belong to different families (according to their chemical characteristics) and are valuable in the food and fuel industries: Lauric acid (fatty acids)(Yunus et al., 2020), squalene (terpenoids)(Englund et al., 2014) and L-leucine (amino acids)(Mustila et al., 2021) (Figure 2). In addition, we examined flux control where biomass was chosen as the objective of analysis, representing a whole-cell product.

**Figure 2.**
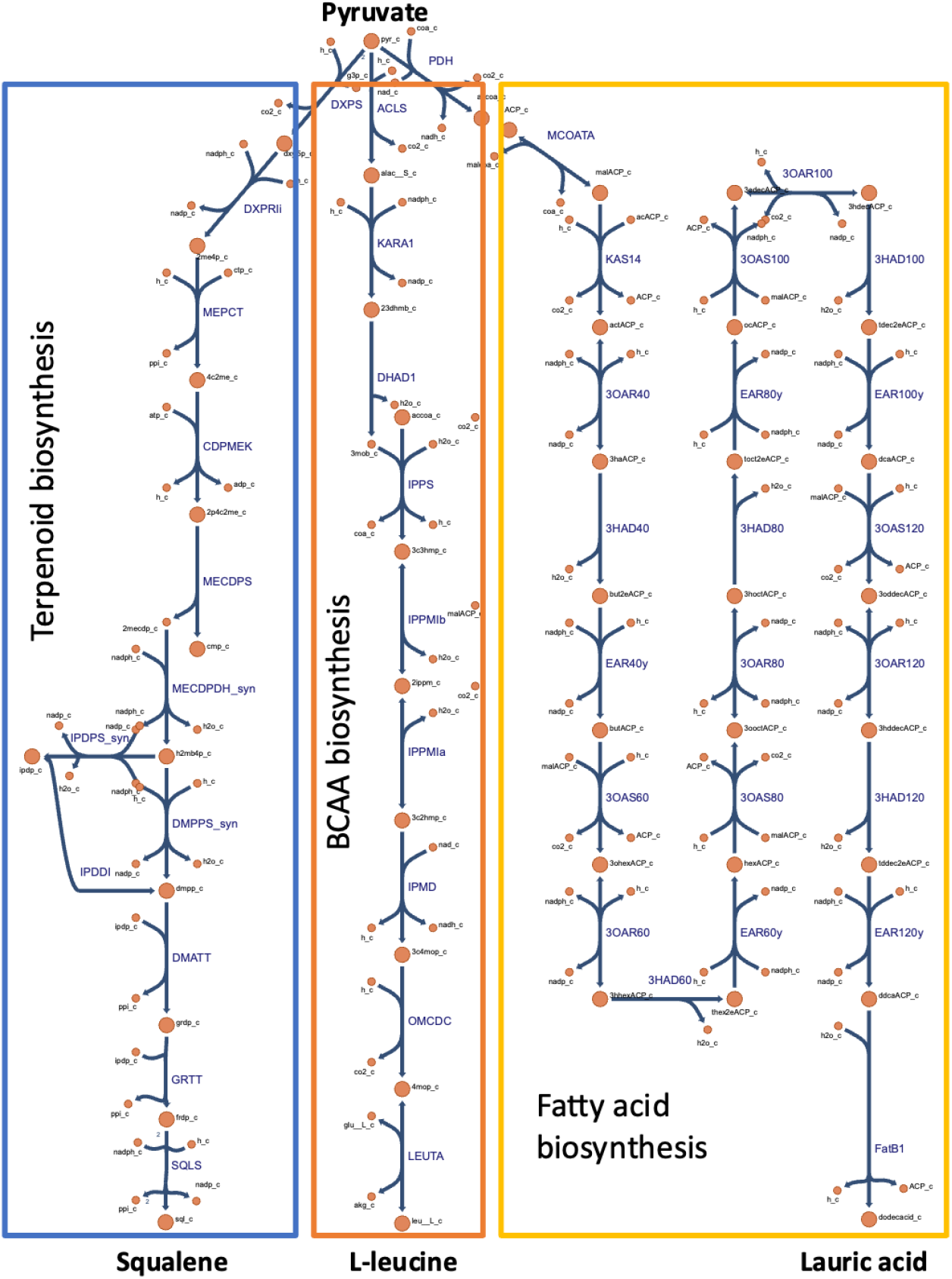
Biosynthesis pathways modelled in the ARCTICA framework for flux control analysis. Metabolic reactions and metabolites, except heterologous ones (FatB1), are indicated by their BiGG identifier (King et al., 2016). BCAA, branched chain amino acids.

### Assessment of ARCTICA performance for discriminating phenotypes

To assess model generalizability and the ability of ARCTICA to classify phenotypes, we examined the predictive metrices. The testing accuracy was 0.99 (Supplementary Table 4) and the F1-score values were 1 (Supplementary Table 5), indicating a high predictive performance of the models for the applied dataset (in silico fluxomic data). The performance of the machine learning model could be also visualized using the confusion matrix (Supplementary Figure 1), where less than 0.1 percent of the trained data was identified as false positive by all the models.

### ARCTICA identifies controlling catalytic steps of the network

To qualitatively evaluate the potential effect of specific reaction rates on the phenotypic outcome, we assessed the resulting distribution of the model coefficients (Figure 3). The analysis shows that the in silico cyanobacterial cell factories are mostly limited by the metabolic capacity of the Calvin–Benson– Bassham (CBB) cycle [ribulose-1,5-bisphosphate carboxylase/oxygenase (RBPC), phosphoribulokinase (PRUK), phosphoglycerate kinase (PGK) and the photosynthetic machinery (PSI_2 and PSII, FNOR) (Figure 3, Figure 4). ARCTICA showed that the enzyme with the highest model coefficient, and thus has the most significant influence on the desired flux (phenotype), was PGK. In comparison, though among the highest influencers, reactions within the product-forming pathway exerted lower flux control over the optimal solution (Figure 2, Figure 3).

**Figure 3.**
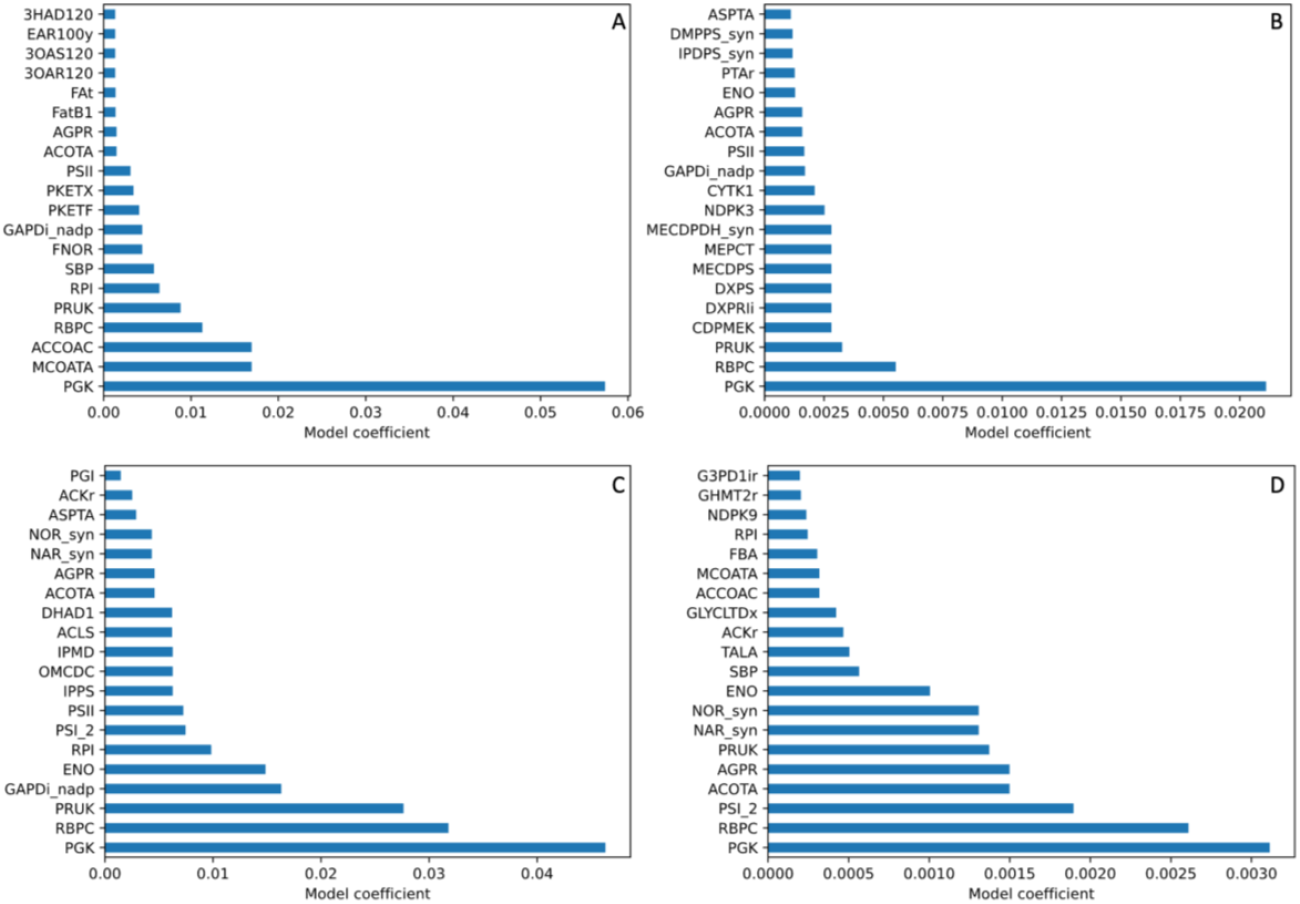
Distribution of model coefficients for the top-20 contributing features (reactions) identified by ARCTICA. A) Lauric acid production. B) Squalene production. C) L-leucine production. D) Biomass. For the full list of metabolic reactions, their corresponding model coefficient and abbreviation, refer to the GitHub repository https://github.com/amitkugler/ARCTICA.

**Figure 4.**
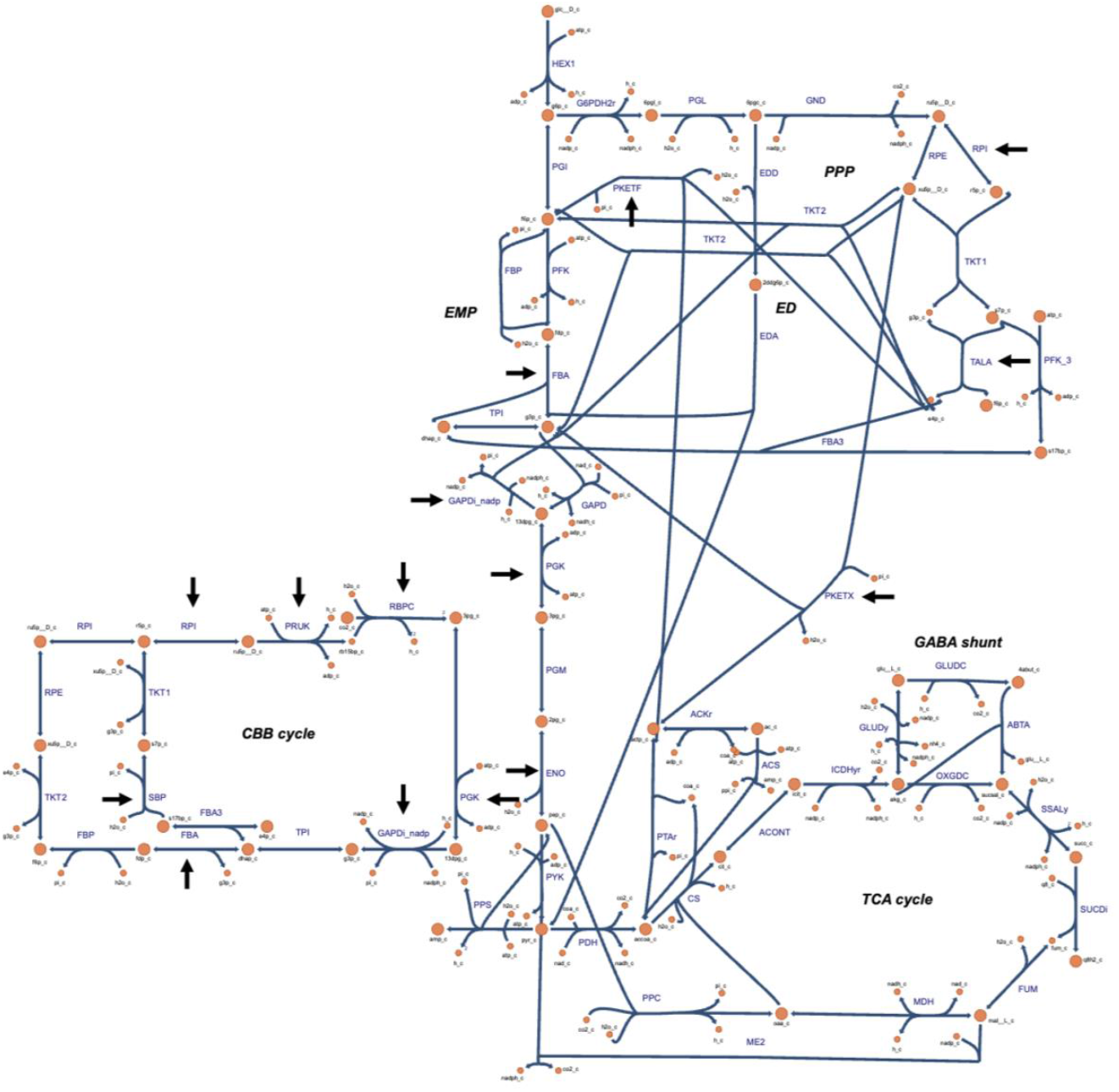
Metabolic map of central carbon metabolism of *Synechocystis* sp. PCC 6803. The map was generated with Escher web-tool (King et al., 2015). Metabolic reactions and metabolites are indicated by their BiGG identifier. TCA, tricarboxylic acid; GABA, branched-chain amino acids γ-aminobutyrate; CBB, Calvin–Benson–Bassham; ED, Entner–Doudoroff; EMP, Embden-Meyerhof-Parnas; PPP, pentose phosphate pathway. Black arrows indicate reactions identified by ARCTICA. For the list of metabolic reactions, their full name and abbreviation, refer to the GitHub repository https://github.com/amitkugler/ARCTICA.

ARCTICA additionally identified product-specific controlling fluxes. For lauric acid production (Figure 3A), acetyl-CoA carboxylase (ACCOAC) and malonyl-CoA-ACP transacylase (MCOATA) were the major determinants of fatty acid synthesis. This was followed by ribose-5-phosphate isomerase (RPI), sedoheptulose-bisphosphatase (SBP) and ferredoxin reductase (FNOR) (Figure 4). The fatty acyl reductase and hydratase enzymes downstream the fatty acid biosynthesis pathway (e.g., EAR100y and 3HAD120) exerted lower limiting effect. For squalene production (Figure 3B), metabolic reactions starting from the pyruvate node until 2-C-methyl-D-erythritol 2,4-cyclodiphosphate were found as high flux-controlling reactions, more than the downstream catalytic enzymes (DMPPS_syn and IPDPS_syn). As a kinetically uncharacterized metabolic pathway, we chose to simulate control points that affect L-leucine biosynthesis (Figure 3C). In addition to the rate of the CBB cycle, ARCTICA found ENO and RPI as flux-controlling reactions (Figure 4), followed by enzymes within the L-leucine biosynthesis pathway, downstream to the 2-ketoisovaleric acid (KIV) node. Further, we were interested in identifying reactions that determine microbial growth, as a whole-cell biomass product (Figure 3D). In a similar manner to production of specific metabolites, we constrained the biomass pseudo-reaction to 10% and 90% of maximal growth rate and applied ARCTICA. We found that after PGK and RBPC, photosystem I activity (PSI_2; ferrocytochrome c oxidation) exerted the most control on biomass formation. A medium-scale control was found for PRUK and ENO, and a weaker control was evaluated for SBP, transaldolase (TALA), fructose-bisphosphate aldolase (FBA) and RPI (Figure 3, Figure 4).

## Discussion

In this study, we showed that by integrating high-throughput fluxomic data with machine learning methods, distribution of flux control can be determined. Thereby, ARCTICA complements previous machine learning-based studies on enzyme parameters (Heckmann et al., 2018; Kroll et al., 2021, 2023; F. Li et al., 2022; Wendering et al., 2023) and extends the traditional MCA. By using substantially fewer parameters and accounting for the nature of modular enzymatic activity within the cell, ARCTICA offers a scalable framework for investigating system-level flux control. Considering model coefficients as indicators for important reactions, we found that, under the conditions examined in this study, our results are in a good agreement with published observations. In order to validate the obtained predictions, we searched for research papers discussing MCA and kinetic measurements. As investigations on *Synechocystis* are scarce, we broadened our comparison to studies performed on other model photosynthetic biological systems, such as *Arabidopsis thaliana*, were applicable.

For example, for all the modelled products, enzymes within the CBB cycle exerted the highest flux control, more than the photosystems. In previous studies, the capability of the CBB cycle appeared to limit electron transport rates in other cyanobacterial species and in plants (Stitt and Schulze, 1994; Stitt, Lunn and Usadel, 2010; Zorz *et al*., 2015; Raines, 2022). Further, ensemble kinetic models identified that PGK exerts a significant control over RuBisCo flux in *Synechocystis* (Janasch et al., 2018; Nishiguchi et al., 2019). Our results are also consistent with previous findings on *Synechocystis*, that the gluconeogenic glyceraldehyde-3-phosphate dehydrogenase (GAPDi_nadp) and enolase (ENO) possess a positive, but weaker, control on the fluxes through the system (Janasch et al., 2018).

ARCTICA, in its exploration of the MCA, revealed bottlenecks in the early stages of fatty acid synthesis. With acetyl-CoA carboxylase (ACCOAC) and malonyl-CoA-ACP transacylase (MCOATA) being contributing the most. While ACCOAC is often considered the key regulator of fatty acid production (Page et al., 1994), its oversynthesis resulted in a disproportionate increase in the malonyl-CoA pool compared to the actual fatty acid synthesis rate (Davis et al., 2000). This suggests that downstream steps, particularly the catalytic rate of MCOATA, impose limitations on fatty acid flux. Studies also indicated that overexpression of the MCOATA biosynthetic gene led to significant fatty acid accumulation in various microalgae species (Chen et al., 2017; Lei et al., 2012; Z. Li et al., 2018), with a strong correlation between MCOATA transcript abundance and fatty acid synthesis levels under stress induction (Tian et al., 2013). ARCTICA identified enoyl-ACP reductase (e.g., EAR100y) and beta-keto-ACP reductase (e.g., 3OAR120) as additional flux-controlling enzymes for lauric acid production, though to a lesser extent than ACCOAC and MCOATA. Earlier independent studies indeed have highlighted a role for both enzymes in the regulation of fatty acid biosynthesis in *E. coli* (Heath & Rock, 1995, 1996; Yu et al., 2011). Furthermore, the promiscuous phosphoketolase (PKETX and PKETF) emerged as an interesting player, providing an alternative and efficient pathway for acetyl-CoA supply (Meile et al., 2001; Xiong et al., 2016). In *S. elongatus* PCC 7942, enhancing the intracellular acetyl-CoA through a heterologous phosphoketolase pathway boosted fatty acid ethyl esters production (Lee et al., 2017). In *E. coli*, a substantial increase in fatty acid production was achieved by overproducing ACCOAC, MCOATA, PGK, and acetyl-CoA-forming enzymes (Xu et al., 2013). Given that the type II fatty acid synthesis (FAS II) pathway in *Synechocystis* is responsible to phospholipid synthesis, similarly to that in *E. coli* (Heath et al., 2002), our findings emphasize the pivotal role of malonyl-CoA-ACP transacylase and phosphoketolase enzymes in the rate of fatty acid production in *Synechocystis*.

Regarding limiting reactions in the methylerythritol 4-phosphate (MEP) pathway, ARCTICA predictions align with previous studies, indicating that the initial steps (DXPS, DXPRIi) hinder isoprenoid production in *Synechocystis* and other photosynthetic organisms (Carretero-Paulet et al., 2006; Englund et al., 2015; Miller et al., 2000; Wright et al., 2014). Notably, DXPS overproduction minimally affected isoprene production in engineered *S. elongatus* PCC 7942 (Gao et al., 2016) and *E. coli* (Lv et al., 2013), while additional overexpression of isopentenyl-diphosphate D-isomerase (IPDDI) biosynthetic gene significantly increased the cyanobacterial isoprene (Englund et al., 2018; Gao et al., 2016) and (E)-α-bisabolene production (Rodrigues & Lindberg, 2021). Our study supports these findings, highlighting IPDDI, as well as 1-hydroxy-2-methyl-2-(E)-butenyl 4-diphosphate reductase (DMPPS_syn), as key regulators of the MEP pathway in *Synechocystis*.

Regarding analysis for L-leucine production, representing an uncharacterized pathway, it was surprising to discover that the reactions converting pyruvate to KIV, [acetolactate synthase (ACLS) and ketol-acid reductoisomerase (KARA1)], exert low control over L-leucine synthesis rate, despite their role in diverting flux from the central carbon metabolism to the branched-chain amino acid biosynthesis pathway.

ARCTICA unveiled both well-known, and novel control points on biomass production. Our focus on enzymes within the CBB cycle, extensively studied in photosynthesis research, identified RBPC, SBP and FBA as key controllers on the overall network flux, aligning with existing experimental and modelling data on cyanobacteria and plants (Janasch et al., 2018; Johnson & Berry, 2021; Poolman et al., 2000; Raines et al., 2008; Zhu et al., 2007). Contrary to prior findings, RPI emerged as a significant control point in our data. SBP, FBA and RPI contribute to ribulose 1,5-bisphosphate (RuBP) regeneration, crucial for the CBB cycle and biomass building block, such as purine nucleotides. SBP and FBA jointly control photosynthetic carbon assimilation with RuBisCO (Raines, 2003), influencing cyanobacterial growth (Liang & Lindblad, 2016). We propose RPI as a potential flux-controlling factor for enhancing bioproduction in *Synechocystis*.

We would like to highlight few points that merit further attention. (1) The machine learning models indicated varying importance levels for predicted reactions, emphasizing the need for a synergistic action of multiple enzymes, rather than relying solely on overproducing individual enzymes (Niederberger et al., 1992; Schaaff et al., 1989). Indeed, the overexpression of *pgk* single gene in *Synechocystis* (Nishiguchi et al., 2019) did not lead to the highest possible production capability of ethanol as estimated by flux balance analysis (Lasry Testa et al., 2019). (2) The control over a metabolic flux is influenced by external conditions (Walsh and Koshland, 1985; Stitt and Schulze, 1994; Nies *et al*., 2023), implying that the identified rate-limiting enzymes may vary under the different growth conditions. (3) The equal importance of product-specific reactions suggests potential overestimation of key enzymes flux capabilities, likely due to uniform sampling of unconstrained flux space and the inability to consider allosteric regulation in this study. Adjustment of ARCTICA using available methods for in vivo kinetic parameters (Davidi et al., 2016; Heckmann et al., 2018) or metabolic fluxes (Hendry et al., 2019), along with constraining the flux space through omics data (Reed, 2012; Sánchez et al., 2017), will enhance the predictive accuracy of the machine learning models.

## Conclusions

We devised a strategy for the identification of controlling biochemical reactions associated with cell phenotype, through statistical evaluation of metabolic features. By considering alternative steady states of intracellular fluxes, we illustrate how cyanobacterial metabolism responds to genetic or environmental perturbations. The proof-of-concept application of ARCTICA demonstrates its capacity to learn mechanistic relations in order to automatically target specific metabolic pathways or enzymes for further functional analysis and strain engineering. While the proposed workflow is computationally intensive, the accessibility of high-performance computers makes this pipeline viable. We envision that this open-source approach will aid computational biologists and experimentalists, and accelerate the creation of efficient cyanobacterial cell factories for diverse chemicals.

## Data availability

The authors declare that all the data supporting the work are available within the paper, its supplementary information, and in the GitHub repository https://github.com/amitkugler/ARCTICA.

## Acknowledgments

This work was supported by Formas—A Swedish Research Council for Sustainable Development” (project no. 2021-01669). The computations were enabled by resources in project [SNIC 2022/22-1100] provided by National Academic Infrastructure for Supercomputing in Sweden (NAISS) at UPPMAX, funded by the Swedish Research Council (project no. 2022-06725). The authors would like to thank Jens Sjölund and Pia Lindberg for valuable discussions.

## CRediT Author contributions

AK: Conceptualization, Methodology, Software, Data curation, Investigation, Formal analysis, Validation, Visualization, Writing-Original Draft, Writing-Review & Editing. KS: Conceptualization, Supervision, Funding Acquisition, Writing-Review & Editing.

