## Supplemental material for "Machine learning predicts system-wide metabolic flux control in cyanobacteria"

**Supplementary material (tables) for Machine learning predicts system-wide flux control in cyanobacteria**

**Supplementary Table 1.** Reactions added to iJN678_AK to create iJN678_AK_dodecanoate.

| **Name** | **Gene–Reaction Rule** | **Description** | **Stoichiometry** | **Reference** |
|---|---|---|---|---|
| FatB1 | UcFatB1 | Acyl-ACP thioesterase | ddcaACP_c + h2o_c --> ACP_c + h_c + dodecacid_c | (Liu et al., 2011) |
| FAt | | Dodecanoate transport via diffusion (cytoplasm to extracellular) | dodecacid_c --> dodecacid_e | |
| EX_FA_e | | Dodecanoate exchange | dodecacid_e--> | |

**Supplementary Table 2.** Reactions added to iJN678 to create iJN678_AK_squalane.

| **Name** | **Gene–Reaction Rule** | **Description** | **Stoichiometry** | **Reference** |
|---|---|---|---|---|
| SQLt | | Squalene transport via diffusion (cytoplasm to extracellular) | sql_c --> sql_e | |
| EX_sql_e | | Squalene exchange | sql_e--> | |

**Supplementary Table 3**. Reactions added to iJN678_AK to create iJN678_AK_leucine.

| **Name** | **Gene–Reaction Rule** | **Description** | **Stoichiometry** | **Reference** |
|---|---|---|---|---|
| LEUt | | Leucine transport via diffusion (cytoplasm to extracellular) | leu__L_c --> leu__L_e | |
| EX_leu__L_e2 | | Leucine exchange | leu__L_e --> | |

**Supplementary Table 4**. Accuracy scores of the machine learning models.

| Objective | Testing accuracy |
|---|---|
| Lauric acid | 1 |
| Squalene | 0.99 |
| L-leucine | 0.99 |
| Biomass | 0.99 |

**Supplementary Table 5**. Classification report of the machine learning models.

| | Precision | Recall | F1-score | Support |
|---|---|---|---|---|
| 1 | 1 | 1 | 1 | 33000 |
| 9 | 1 | 1 | 1 | 33000 |
| accuracy | | | 1 | 66000 |
| macro avg | 1 | 1 | 1 | 66000 |
| weighted avg | 1 | 1 | 1 | 66000 |

**Supplementary material (figures) for Machine learning predicts system-wide flux control in cyanobacteria**

Amit Kugler[a], Karin Stensjö[a]*

[a]Microbial Chemistry, Department of Chemistry-Ångström Laboratory, Uppsala University, Box 523, SE-751 20, Uppsala, Sweden

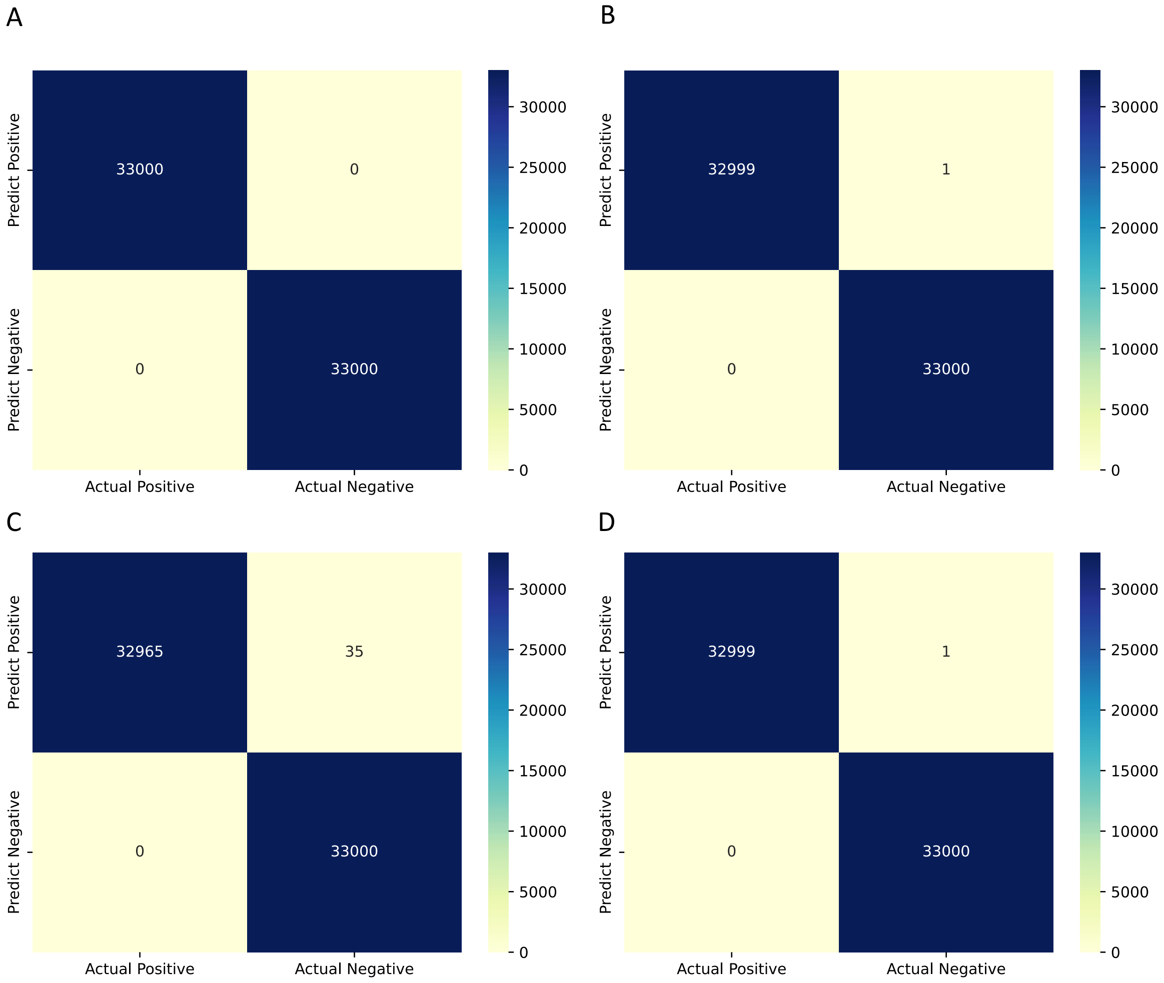

**Supplementary Figure. 1** Features selection by logistic regression (LASSO regularization). Resulting confusion matrix based on the machine learning models.
